# Metagenome-assembled genomes provide insight into the microbial taxonomy and ecology of the Buhera Soda Pans, Zimbabwe

**DOI:** 10.1101/2024.02.15.580475

**Authors:** Ngonidzashe Mangoma, Nerve Zhou, Thembekile Ncube

## Abstract

The use of metagenomics has substantially improved our understanding of the taxonomy, phylogeny and ecology of extreme environment microbiomes. Advances in bioinformatics now permit the reconstruction of almost intact microbial genomes, called metagenome-assembled genomes (MAGs), from metagenomic sequence data, allowing for more precise cell-level taxonomic, phylogenetic and functional profiling of uncultured extremophiles. Here, we report on the recovery and characterisation of metagenome-assembled genomes from the Buhera soda pans located in eastern Zimbabwe. This ecosystem has not been studied despite its unique geochemistry and potential as a habitat for unique microorganisms. Metagenomic DNA from the soda pan was sequenced using the DNA Nanoball Sequencing (DNBSEQ^R^) technique. Sequence analysis, done on the Knowledgebase (KBase) platform, involved quality assessment, read assembly, contig binning, and MAG extraction. The MAGs were subjected to taxonomic placement, phylogenetic profiling and functional annotation in order to establish their possible ecological roles in the soda pan ecosystem. A total of 16 bacterial MAGs of medium to high quality were recovered, all distributed among five phyla dominated by *Proteobacteria* and *Firmicutes*. Of the ten MAGs that were taxonomically classified up to genus level, five of them belonged to the halophilic/ haloalkaliphilic genera *Alkalibacterium, Vibrio, Thioalkalivibrio, Cecembia* and *Nitrincola*. Functional profiling revealed the use of diverse carbohydrate-metabolising pathways among the MAGs, with glycolysis and the pentose phosphate pathways appearing to be key pathways in this ecosystem. Several MAGs harboured both sulphur/ sulphate reduction and respiratory pathways, suggesting a possible mechanism of energy generation through sulphur/ suphate respiration. In conclusion, this study revealed a highly taxonomically and functionally diverse microbial community in the soda pans, dominated by halophilic and haloalkaliphilic bacteria.

## 1. Introduction

The Buhera soda pans are a unique, naturally occurring saline and alkaline aquatic system located in a remote part of the Buhera district found in Eastern Zimbabwe. No record exists at present regarding the scientific exploration of this potentially important extreme environment. However, in our recent exploration of this site, we established that the Buhera soda pans are highly alkaline (pH 8.74 – 11.03), moderately saline (2.26 – 2.94 mg/l) and belong to the carbonate type of soda pans and lakes (Mangoma *et al*., unpublished). Our findings are consistent with several studies that reported on the geochemistry of various soda pans and The geochemistry of the Buhera soda pans matches that of several other soda pans and lakes from around the world (He *et al*., 2022; Felföldi, 2020; Sorokin *et al*., 2018). In general, soda pans and lakes have alkaline pH in the range of 8 – 11, and highly variable salinity with a lower threshold of around 1 mg/l (Boros and Kolpakova, 2018). The extreme and distinctive geochemistry of the Buhera soda pans exerts selection pressure on their microbiomes, often leading to the evolution of unique microbial communities. The composition of the microbial communities of soda pans and lakes, and the possibility of finding novel microorganisms with tangible biotechnological potential often attracts interest in these microbial habitats.

The extreme conditions existent in soda pans and lakes modify their microbial communities, leading to the dominance of extremophiles, with halophiles, alkaliphiles and haloalkaliphiles often dominating these microbiomes (Bryanskaya *et al*., 2022). The ability of different microbial taxa to thrive under the alkaline and saline conditions of soda pans stimulate interest among microbiologists about such microbial ecosystems, and elevate the possibility that some of the microorganisms that inhabit these environments have potentially useful applications. For instance, numerous studies have demonstrated that soda pans and lakes harbour a rich diversity of novel microbes, enzymes and unique microbial metabolites (Sorokin *et al*., 2018). Enzymes such as amylases, proteases, lipases and cellulases have been extracted from haloalkaliphiles isolated from saline-alkaline systems such as soda pans, and some of these enzymes have been demonstrated to perform optimally under conditions of alkaline pH and salinity (Nyakeri *et al*., 2018). Enzymes that retain their stability and high performance under alkaline pH and salinity have several potential applications in industrial processes such as leather tanning, starch saccharification and other food processes, detergent making and the bioremediation of saline and/ or alkaline effluents such as those from the fat/ oil and soap industry. The potential benefits from the exploration of the microbial communities of soda pans and lakes provide impetus for more inquiry into their microbial communities. A combination of both culture-based and culture-independent techniques have helped expose the rich microbial communities of soda pans and lakes (Choure *et al*., 2021; Govil *et al*., 2021; Sysoev *et al*., 2021; Guan *et al*., 2020). However, it is estimated that up to 99% of the microbial inhabitants of extreme environments are unculturable (Jin *et al*., 2019; Grötzinger *et al*., 2018), leading to an increasing reliance on techniques such as metagenomics in studies of extreme microbiomes.

Recent innovations in DNA extraction, sequencing and sequence analysis technologies have exponentially broadened the scope of metagenomic studies of extreme environments, leading to the exploration of many new extreme environments and the discovery of new microbial taxa. While most metagenomics studies allow for read-based community-level determination of microbial community composition and function, new developments now make it possible to perform more precise cell-level metagenomic analysis using techniques such as metagenome-assembled genome (MAG) construction. A metagenome-assembled genome (MAG) is a hypothetical microbial genome created using contigs derived from the assembly of metagenomic sequence reads (Chivian *et al*., 2023). Consequently, each MAG represents a putative microbial genome, and may contain enough information to enable its taxonomic placement, phylogenetic profiling and functional annotation (Singh *et al*., 2023). MAG extraction is an innovative technique that allows not only for metagenomic sequence analysis to be performed at the level of individual microbes, allowing more accurate description of their taxonomy and ecological function, it also allows for the discovery and characterisation of previously uncultured microorganisms especially from extreme environments such as soda pans. Thus, being able to recreate MAGs from extreme environment metagenomic DNA provides a back-door channel through which we can explore the total microbial community of an extreme environment, with the usual limitations imposed by the physico-chemical demands of extremophiles.

In this study, we report on efforts to construct and characterise metagenome-assembled genomes (MAGs) from metagenomic DNA obtained from the Buhera soda pans. Several parameters are used to characterise the integrity of a MAG, and one such key parameter is MAG completeness, which indirectly reflects the size of the MAG relative to the reference genome. While many studies report on MAGs with completeness in the 50 – 60% range (Singh *et al*., 2023), we report on the creation of several MAGs with completeness above 80%. The high degree of completeness enables one to then make a variety of highly accurate predictions about an organism’s taxonomy, phylogeny and physiology, making it possible to precisely allocate ecological functions to uncultured extremophiles.

## 2. Materials and Methods

### 2.1 Metagenomic sequence data

The metagenomic sequence data used in this study was obtained from a previous study on the Buhera soda pans. The metagenomic DNA used was extracted using the ZymoBIOMICS DNA Miniprep kit (Zymo Research, CA, USA), and sequenced using the DNA nanoball sequencing (DNBSEQR) technique by BGI Genomics company, Hong Kong.

### 2.2 Metagenomic sequence analysis

Metagenomic sequence analysis was performed on the United States Department of Energy (DOE) Systems Biology Knowledgebase (KBase) pipeline (Arkin *et al*., 2018), an integrated sequence analysis platform with several in-built functions that allow for the execution of a number of sequence analysis tasks such as quality assessment, sequence trimming, sequence assembly, binning, and metagenome-assembled genome (MAG) extraction and annotation (Figure 1).

**Figure 1:**
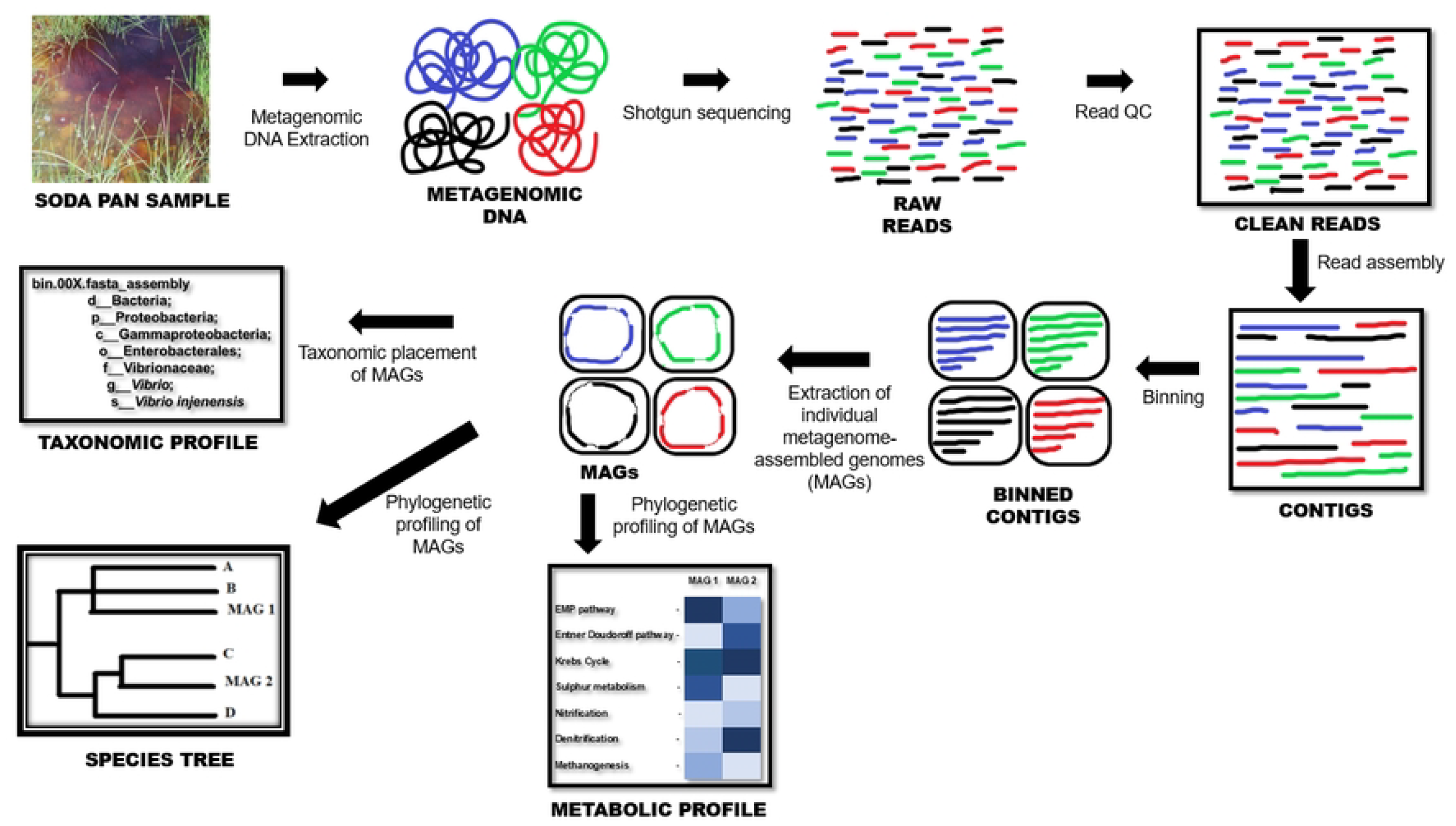
Overview of the sequence analysis workflow employed in this study.

#### 2.2.1 Read assembly

Clean, QC-checked metagenomic sequence reads were assembled into contigs using the two de-novo sequence assemblers SPAdes (v3.15.3) and hybridSPAdes (v3.15.3), with other parameters for both assemblers set at default and minimum contig length of 500. The two assemblers perform contig construction based on sequence *k-mer* composition and make use of *de Bruijn* graphs (Antipov *et al*., 2016; Bankevich *et al*., 2012). The contig libraries generated were compared in KBase using the “Compare Assembled Contig Distributions” software (v1.1.2), an application that compares contig libraries according to parameters such as number of contigs, maximum contig length, contig length distribution and average contig length. This comparison showed that the hybridSPAdes-generated contig library had better features and was used for the subsequent binning exercise.

#### 2.2.2 Contig binning

The hybridSPAdes-generated contigs were binned using the three binning tools: MaxBin2 (v2.2.4), MetaBat2 (v1.7) and CONCOCT (v1.1). MaxBin2 was set at default, with the following additional parameter settings: prob_threshold – 0.8; marker set – 107 bacterial marker gene set; minimum contig length – 1000. MetaBat2 was run with all parameters set at default, with minimum contig length of 2500. Finally, parameters for the CONCOCT binning application were set as follows: read mapping tool – Bowtie 2; minimum contig length – 2500; contig split size – 10 000; *kmer* length – 4; maximum number of clusters for VGMM – 400; maximum number of iterations for VGMM – 500. The resultant bin libraries were compared for such features as bin coverage and number of bins generated. The three bin libraries were then optimised using the DAS-Tool (v1.1.2) at default setting, using diamond as the gene identification tool. The DAS tool enables one to generate optimum consensus high-quality bins using, as input, the binning outputs of more than one binning app. In this study, DAS would examine the bins created by the MaxBin2, MetaBat2 and CONCOCT binning tools and come up with a consensus library of high-quality bins from the three bin libraries. Subseuently, the three original bin libraries (MaxBin2, MetaBat2 and CONCOCT) and the DAS tool-generated bin library were compared for coverage and quality and it was noted that the DAS tool-generated bin library had better features, leading to its selection and use for all downstream steps.

#### 2.2.3 Metagenome-Assembled Genome (MAG) extraction and annotation

The DAS tool-generated bin library was firstly checked for completeness and contamination using the CheckM tool (v1.0.18), using default settings. Thereafter, metagenome-assembled genomes (MAGs) were extracted from the DAS bin library using the “Extract Bins as Assemblies from Binned Contigs” application (v1.0.2), at default settings. The extracted MAGs were annotated using the RASTtk (v1.073), using default settings. The MAG annotation involved determining the size of the genome as well as the detection, identification and functional annotation of open reading frames (ORFs) and genome features such as genes coding for rRNA, tRNA, selenoproteins and pyrrolysoproteins, and clustered regularly interspaced short palindromic repeat (CRISPR) regions present in the MAGs. Additionally, whole-genome maps for selected MAGs were created and annotated using the Proksee web-based application. Proksee allows one to create a complete circular or linear map of bacterial (or other) genomes, and comes with several integrated tools that allow for detailed annotation of the genome maps (Grant *et al*., 2023). In this study, the MAGs were annotated using the following tools: Comprehensive Antibiotic Resistance Database (CARD) Resistance Gene Identifier (RGI) for the identification of antibiotic resistance genes (v1.2.0) (Alcock *et al*., 2023), CRISPR/ CasFinder (v4.2.20) for the identification of CRISPR/ Cas regions (Couvin *et al*., 2018), and Prokka (v1.14.6) for the annotation of coding sequences and RNA regions (Seemann, 2014).

#### 2.2.4 Taxonomic and phylogenetic profiling of MAGs

Annotated MAGs were taxonomically classified using the GTDB-Tk taxonomic classification tool (v1.7.0) using default settings and the following additional settings: minimum alignment percentage – 10; genetic code – 11 (for archaea, most bacteria, most viruses and some mitochondria). The GTDB-Tk tool performs taxonomic classification of the MAGs using single-copy phylogenetic markers to place each genome (MAG) into the GTDB species tree (https://narrative.kbase.us/#catalog/apps/kb_gtdbtk/run_kb_gtdbtk). The MAGs were also inserted into species trees using the “Insert Set of Genomes into Species Tree” app (v2.2.0). This app shows the phylogenetic relationships among the MAGs as well as with other microbes archived in different reference databases.

#### 2.2.5 MAG functional profiling

Apart from taxonomic placement, the MAGs were also subjected to functional profiling using the “Annotate and Distill Assemblies with DRAM” (v0.1.2) application using default settings and the following additional parameters: minimum contig length – 2500; translation table – 11; bit score threshold – 60; reverse search bit score threshold – 350. The DRAM tool searches the MAGs for functional marker genes and groups them into pathways. The pathway completeness for each MAG is determined, making it possible to ascertain likely metabolic roles of each MAG in the microbial community. In this study, DRAM was used to determine pathway completeness for pathways involved in carbon metabolism, nitrogen and sulphur metabolism, electron transport systems and other vital cellular functions. Additionally, more specific gene family annotation of the MAGs based on Hidden Markov Models (HMMs) was performed using the HMMER program. In this case the dbCAN collection of HMMs built from the Carbohydrate-Active Enzymes (CAZy) was determined using HMMER (v10) at default settings.

## 3. Results and Discussion

### 3.1 High-quality metagenomic sequence reads were obtained

The metagenomic DNA used in this study was obtained from another ongoing unpublished study of the Buhera soda pans. The sequence obtained from the DNBSEQ^R^ was of high quality as shown in Table 1:

**Table 1:** Summary of Buhera soda pan metagenomic sequence file.

| Object Name | Buhera Soda Pan Metagenomic Sequence |
| --- | --- |
| Number of Reads | 35,876,873 |
| Total Number of Bases | 3,587,687,300 |
| Mean Read Length | 100.0 |
| Base quality score (range) | 34 – 36 |
| Read quality score (range) | 24 – 38 |
| GC Percentage | 52.08% |

As shown in Table 1, the sequences obtained were composed of 35.9 million short reads averaging 100 bp in size each. The reads were of high quality, with read quality scores in the range of 24 – 38.

### 3.2 HybridSPAdes assembler produced most optimum contig library

Two distinct contig libraries were created using the two de-novo assemblers, SPAdes and hybridSPAdes. However, upon comparison for key contig library features such as contig number, total assembled sequence length, N50 and L50, the hybridSPAdes contig library appeared to have better features (Table 2).

**Table 2:** Features of the SPAdes and hybrid SPAdes assembly libraries. The values highlighted represent the best output for each parameter.

| Contig library feature | ASSEMBLY LIBRARY |  |
| --- | --- | --- |
|  | hybridSPAdes.Assembly | SPAdes.Assembly |
| Number of contigs | 104.964 | **<u>100,265</u>** |
| Number of contigs ($\geq 10000$ bp) | **<u>1,046</u>** | 1,030 |
| Number of contigs ($\geq 100000$ bp) | **<u>55</u>** | 53 |
| Largest contig (bp) | **<u>312,583</u>** | **<u>312,583</u>** |
| Total length (bp) | **<u>145,298,474</u>** | 136,513,900 |
| Total length ($\geq 1000$ bp) | **<u>96,753,868</u>** | 89,478,069 |
| Total length ($\geq 10000$ bp) | **<u>33,554,519</u>** | 32,804,010 |
| Total length ($\geq 100000$ bp) | **<u>8,156,189</u>** | 7,789,317 |
| N50 | **<u>1697</u>** | 1675 |
| L50 | 13884 | **<u>12822</u>** |

Read assembly entails combining short sequence reads produced by high-throughput sequencing platforms in order to recreate the longer fragments, called contigs. Assembly is often the first post-QC step in metagenomic sequence analysis. However, the success of read assembly depends on factors such as the purity of the DNA molecules sequenced, the quality of sequencing output, and sequencing depth. During sequencing, individual reads should sufficiently overlap in a way that makes it possible to form longer consensus sequences during assembly (Alneberg, 2018). In this study, the hybridSPAdes assembler managed to assemble a larger proportion of available reads, producing a larger number of contigs greater than 100 kb when compared to SPAdes. In general, a good assembler should produce a small number of large contigs, covering as much of the available sequence reads as possible, and should produce a contig library with high N50 (the length of the shortest contig at 50% of the total assembly when contigs are arranged in descending order of size) and low L50 (the number of contigs, starting with the longest contig, required to create 50% of the total assembly size) values. Based on this broad criteria, hybridSPAdes was adjudged to have been relatively superior, and hence the hybridSPAdes library was used for the subsequent binning step.

### 3.3 Contig binning and bin optimisation

The three binning tools used produced different amounts of bins, with CONCOCT producing the most bins at 37, followed by 31 produced by MetaBAT2 and lastly MaxBin2 which produced only 25 bins. Each bin is a collection of contigs that have significant homology and, therefore, most likely come from the same genome. While the three binning tools produced different numbers of bins, further analysis revealed that MaxBin2 managed to place a larger proportion of available contigs (24.81%) into bins as compared to the CONCOCT (6.56%) and MetaBAT2 (5.21%) tools. The ability of the MaxBin2 tool to place more contigs into bins will likely produce bins with a higher degree of completeness, which usually leads to the generation of higher quality MAGs. Another key observation is that even though MaxBin2 managed to place only 24.81% of the 100 265 contigs available into bins, the total length of these binned contigs amounts to more than half (55.4%) of the assembled sequence length. This further highlights the superiority of the MaxBin2 binning tool in the specific context of this study. The MaxBin2 bin library was therefore determined to be the best of the three bin libraries. Additionally, bin optimisation using the DAS tool created a consensus bin library composed of 16 bins using, as input, the output of the MaxBin2, MetaBAT2 and CONCOCT tools. A comparison of these 4 bin libraries clearly showed the superiority of the DAS bin library over the other three (Figure 2).

**Figure 2:**
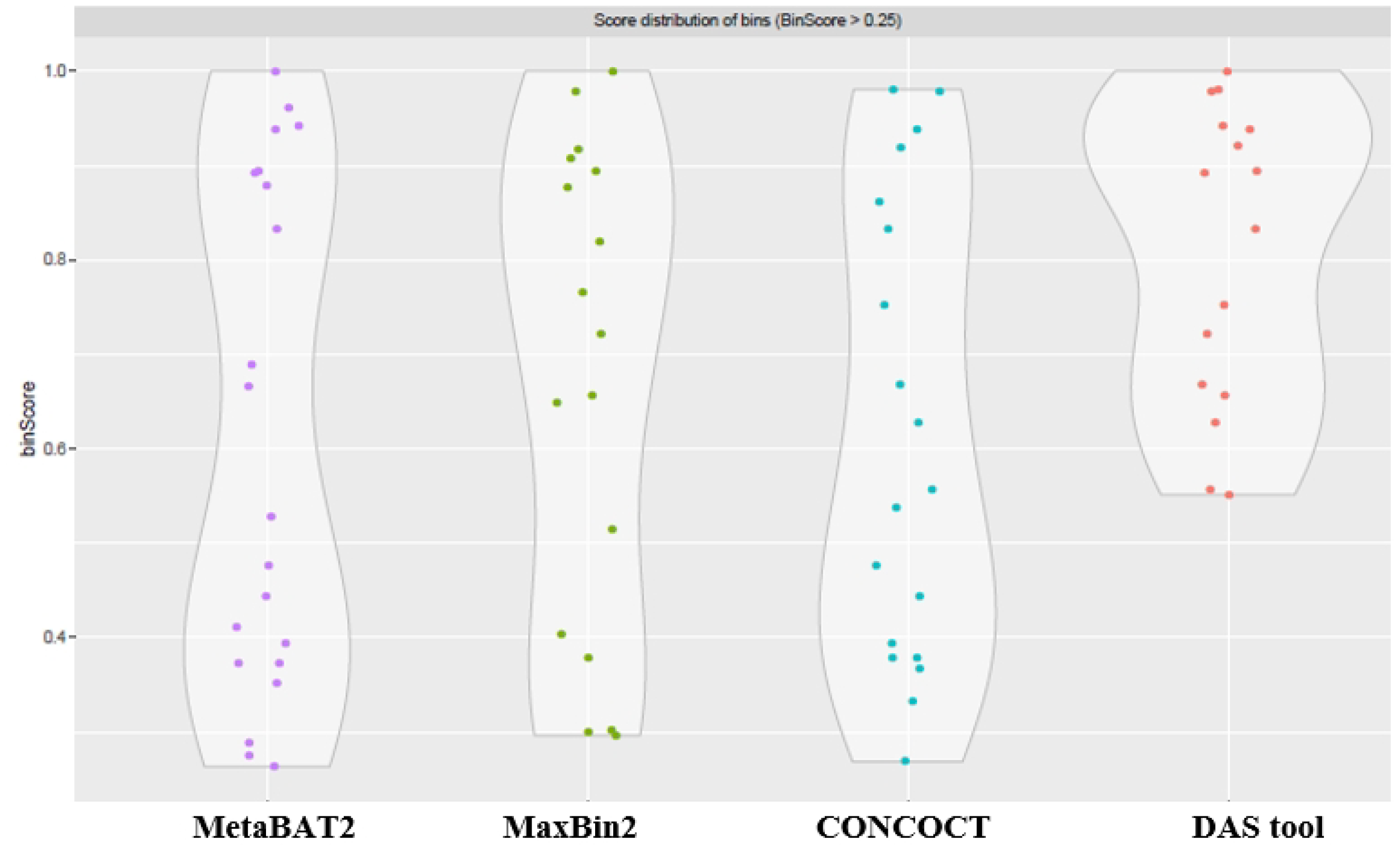
Comparison of the four different bin libraries created in this study. The different coloured dots represent individual bins. Bin library comparison is based on bin score, and only bins with a bin score greater than 0.25 are shown.

As shown in Figure 2, the DAS bin optimisation tool assembled a library of bins of relatively higher quality by consolidating bins from the other binning apps. In this case, DAS selected only bins with a minimum bin score of 0.5. The DAS bin library represents a consolidation of high-quality bins from the other binning app, and as a consequence was used for MAG extraction.

### 3.4 Sixteen medium-to-high-quality MAGs were recovered from the Buhera soda pans

Before their annotation, the MAGs were checked for quality and a majority of them were found to have high levels of completion and low levels of completeness (Figure 3).

**Figure 3:**
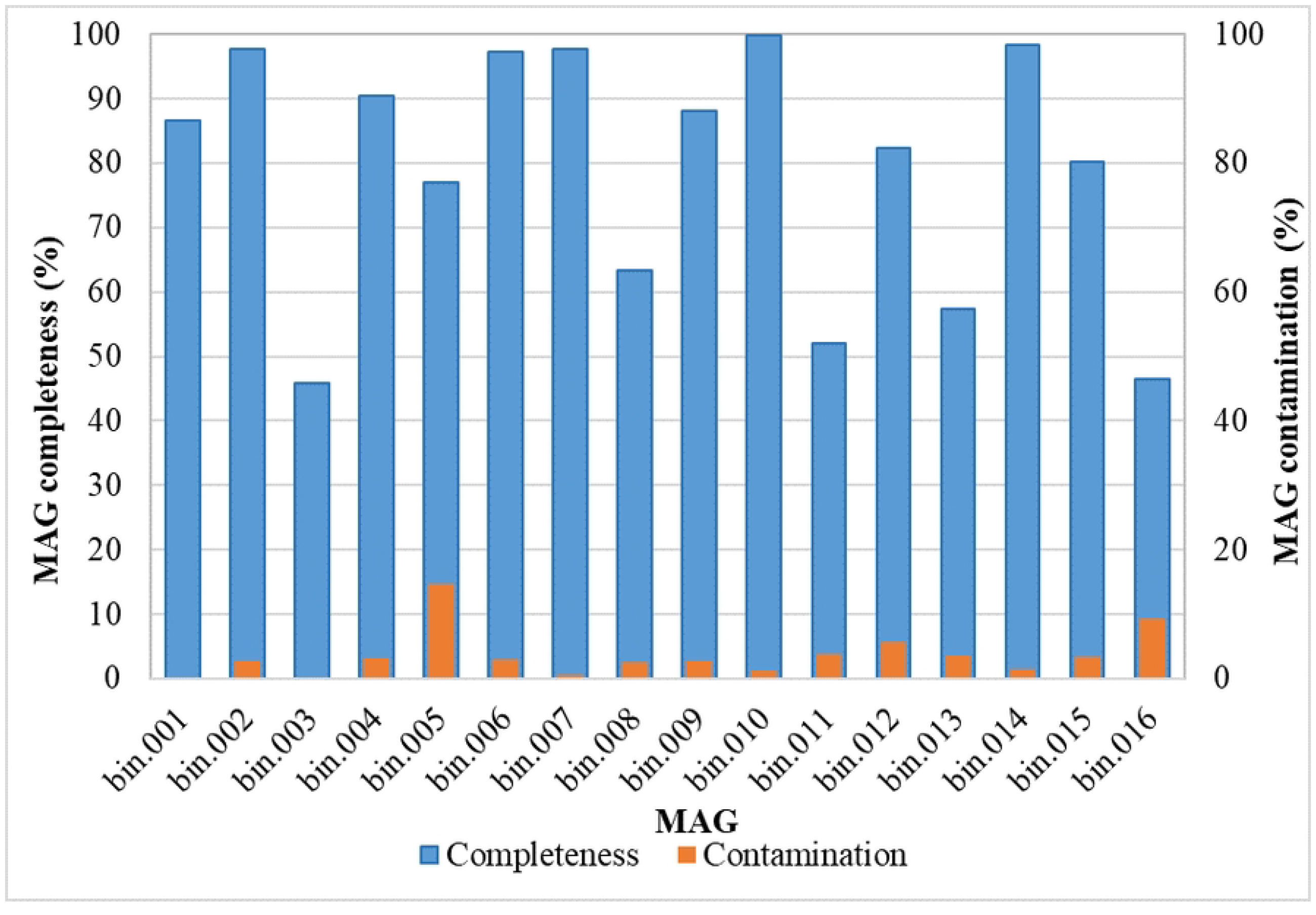
CheckM quality assessment of MAG quality.

MAG completeness and level of contamination are important measures of MAG quality. MAG completeness refers to the fraction of a microbial genome captured in a MAG. The level of completeness of a MAG is important in determining the type of downstream analysis possible, for instance, complete genomes are desired for downstream analyses such as pangenome analysis and genetic linkage studies, whereas partial genomes may be suitable for metabolic prediction and phylogenetic profiling (Bowers *et al*., 2017). The CheckM tool (v1.0.18) used in this study to estimate the level of contamination of the MAGs uses unique, single-copy, universal marker genes found in nearly all taxa to estimate completeness (Bowers *et al*., 2017). On the other hand, the fraction of “foreign” sequences contained in a MAG determines its level of contamination. Consequently, MAGs may be classified, based on their levels of completeness and contamination, as finished, high-quality, medium-quality or low-quality drafts. Under this classification, “finished” MAGs should be of the highest quality, being composed of a single uninterrupted sequence, while “high-quality draft” MAGs are characterised by a minimum completeness of 95% and contamination of less than 5%, whilst encoding the 23S, 16S, and 5S rRNA genes, as well as tRNAs for at least 18 of the 20 possible amino acids (Bowers *et al*., 2017). Lastly, MAGs with at least 50% contamination and less than 10% contamination are considered “medium-quality draft” MAGs, while “low-quality draft” MAGs have less than 50% completeness and less than 10% contamination.

On the basis of this classification, 31% of the 16 MAGs produced were classified as “high-quality draft” MAGs, 63% were “medium-quality draft” MAGs, whereas only 1 MAG (bin 0.03) was of low quality with a completeness of 45.95%. The medium-and high-quality MAGs obtained in this study meet the minimum requirements for “Minimum Information on Metagenome-assembled Genomes” (MIMAGs) as proposed by the Genomics Standards Consortium (Bowers *et al*., 2017). Overall, the quality of most MAGs obtained in this study is suitable for further downstream analysis of the genomes obtained. Having high quality MAGs is advantageous in that the MAGs created tend to highly resemble actual bacterial genomes, making subsequent analyses such as taxonomic placement, functional profiling, and metabolic modelling easier (Chivian *et al*., 2023). In fact, MAG extraction is an attempt at reconstructing microbial genomes from metagenomic sequences. A MAG is a highly valuable tool as it enables a researcher to do more than just taxonomic classification of organisms, which is often the case with read-based analysis. A high-quality MAG will possess most of the genetic elements of a typical microbial genome, such as phylogenetic marker genes, protein-coding genes and regulatory genes, making it easier to derive more insight on the ‘hypothetical’ microbe which it represents in the context of its taxonomy and ecological role in the environment. MAG creation also makes it possible to uncover new and previously uncultured microorganisms by firstly rebuilding their genomes. The first bacterial genomes to be reconstructed from metagenomic data originated in samples from acid mine drainage water (Anelberg, 2018).

### 3.5 Annotation reveals structural and organizational diversity of recovered genomes

The 16 MAGs obtained in this study were highly diverse in terms of size and organisation, as exemplified by the differential distribution of genome features such as coding and non-coding genes, non-coding repeats and RNA-coding genes (Figure 4).

**Figure 4:**
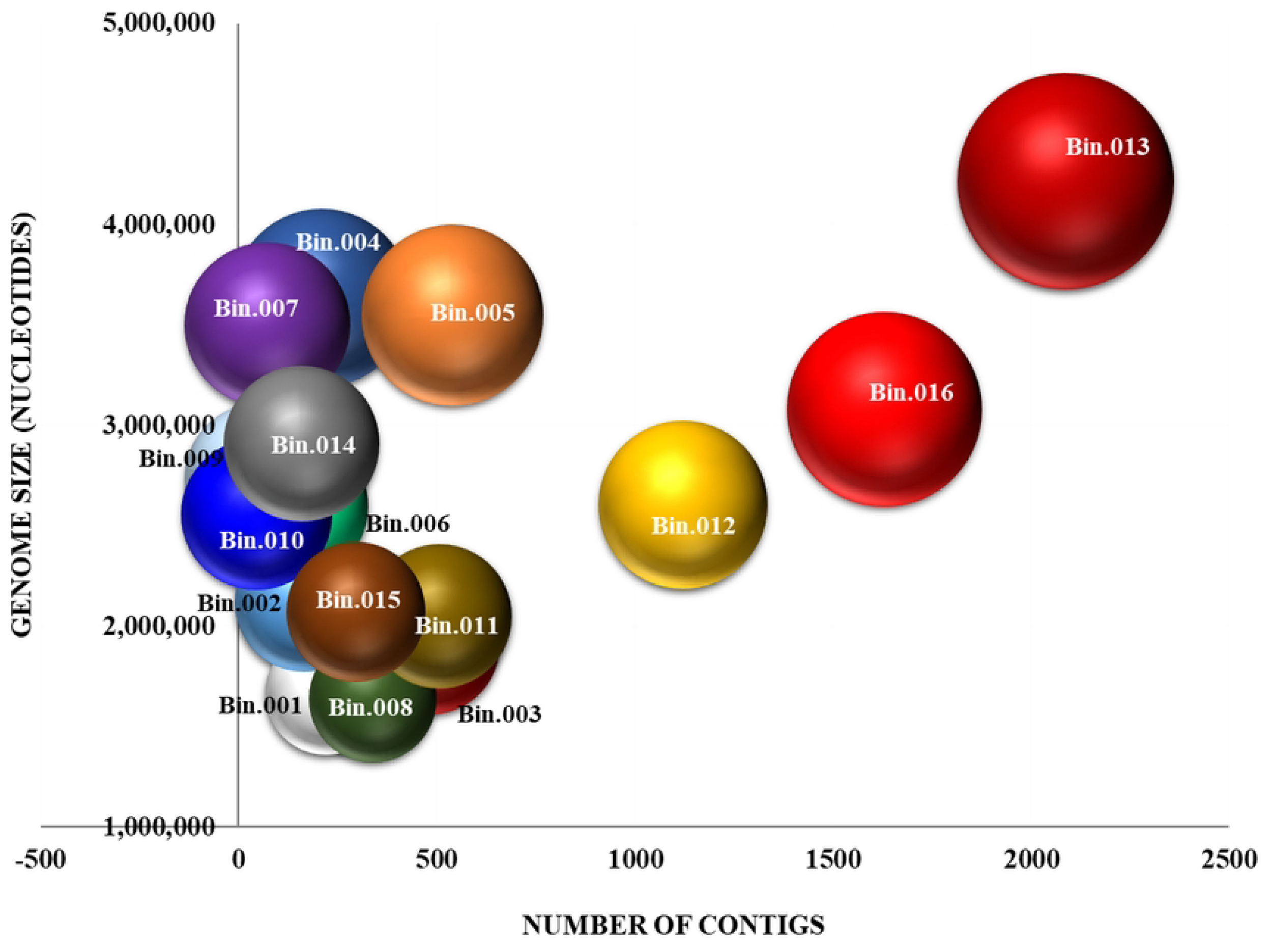
Annotation of MAGs using the RASTtk application. Each bubble represents a MAG. The position of a bubble shows the number of contigs making up the MAG (horizontal axis) and the size of the MAG (in nucleotides). The size of each bubble is proportional to the number of genome features detected in the MAG.

The largest MAG created in this study (Bin.013) was 4,415,129 bp in size. Four other MAGs (Bins 004, 005, 007 and 016) exceeded 3 Mb in size. These MAGs are within range of the size of the average bacterial genome. Though bacterial genome size varies considerably, it is generally accepted that the average bacterial genome is about 5 Mb in size and encodes approximately 5000 proteins (Land *et al*., 2015). The closer a MAG is to an actual genome in terms of size, the more information one will get of its genetic elements and metabolic potential upon analysis. This enables one to get more insight about the microbial inhabitants of an environment and their possible ecological roles in the ecosystem. The annotation tool used to annotate genome features of the MAGs in this study, RASTtk, calls open reading frames using both Prodigal (Hyatt *et al*., 2010) and Glimmer3 (Delcher *et al*., 2007). The genome features were functionally annotated using the algorithms Kmers V2, Kmers V1 and protein similarity. RAST uses the k-mer approach to find genes in reference databases that are homologous to the query sequences, using a collection of gene family functional annotations to rapidly annotate any matches (Chivian *et al*., 2023).

### 3.6 All recovered MAGs belong to only five phyla under domain bacteria

All the MAGs recovered from the Buhera soda pans were shown to belong to domain bacteria, all being distributed among the five phyla *Proteobacteria, Firmicutes, Chloroflexi, Bacteroides* and *Deinococcus* (Table 3).

**Table 3:**
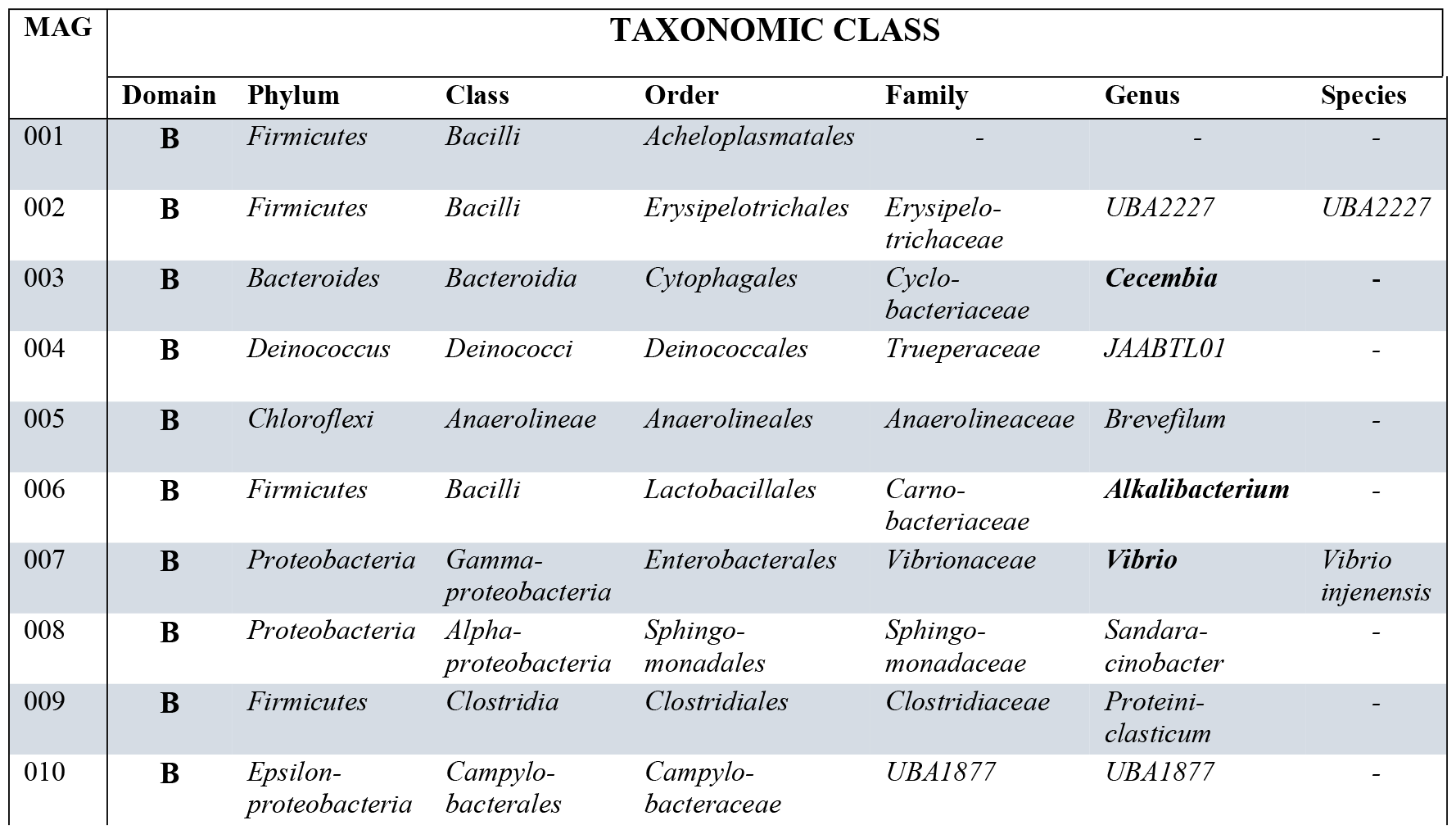
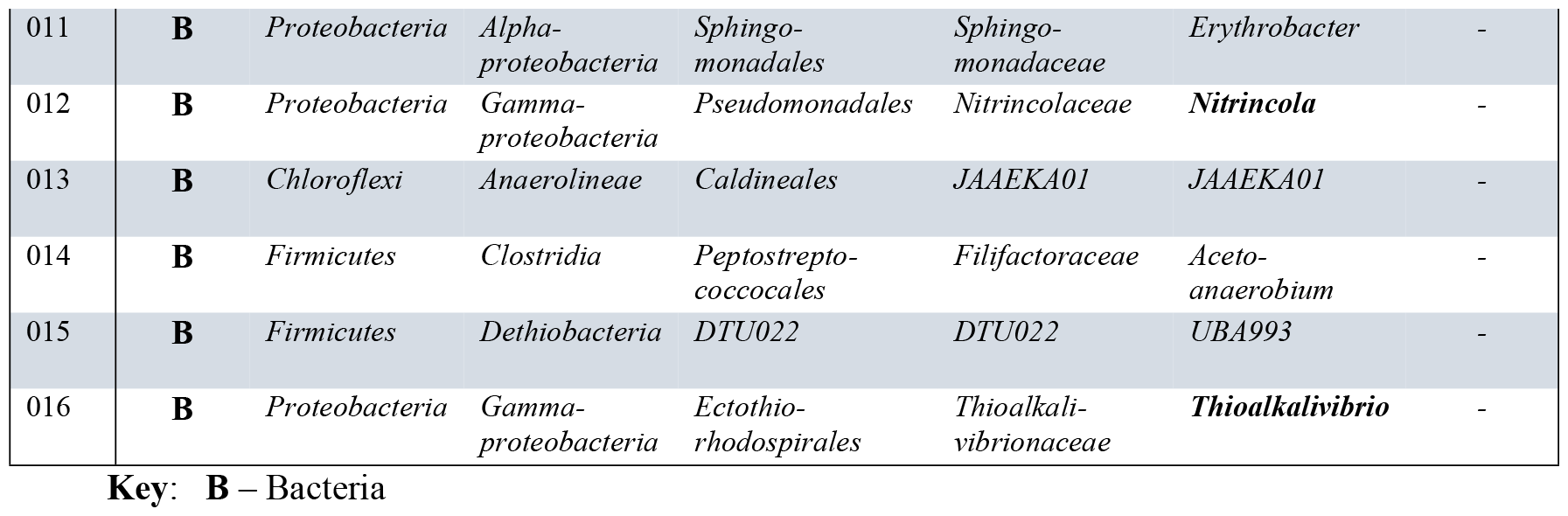
Taxonomic placement of MAGs. The taxonomic classification as well as the identity of closest match from reference databases, where available, is shown.

The two bacterial phyla, *Proteobacteria* and *Firmicutes*, accounted for 75% of all MAGs created, asserting their dominance in the Buhera soda pans. This observation is consistent with our findings in an earlier study where read-based taxonomic analysis showed that *Proteobacteria* and *Firmicutes* constituted 67% of the Buhera soda pan microbiome (Mangoma *et al*., UNPUBLISHED). Several studies have also reported on the dominance of phylum *Proteobacterium* and *Firmicutes* in soda pans and lakes (He *et al*., 2022; Omeroglu *et al*., 2021; Felföldi, 2020)). Additionally, ten out of the sixteen MAGs were classified up to genus level, with a notable finding being the presence of the following halophilic/ haloalkaliphilic genera: *Alkalibacterium, Vibrio, Thioalkalivibrio, Cecembia* and *Nitrincola* being. These halophiles/ haloalkaliphilic genera are commonly associated with saline and saline-alkaline environments, thus their presence in abundance in the Buhera soda pans highlights its status as a natural habitat for salt and alkali-loving extremophiles.

Additionally, all MAGs were placed into individual species trees as shown in Figure 5.

**Figure 5:**
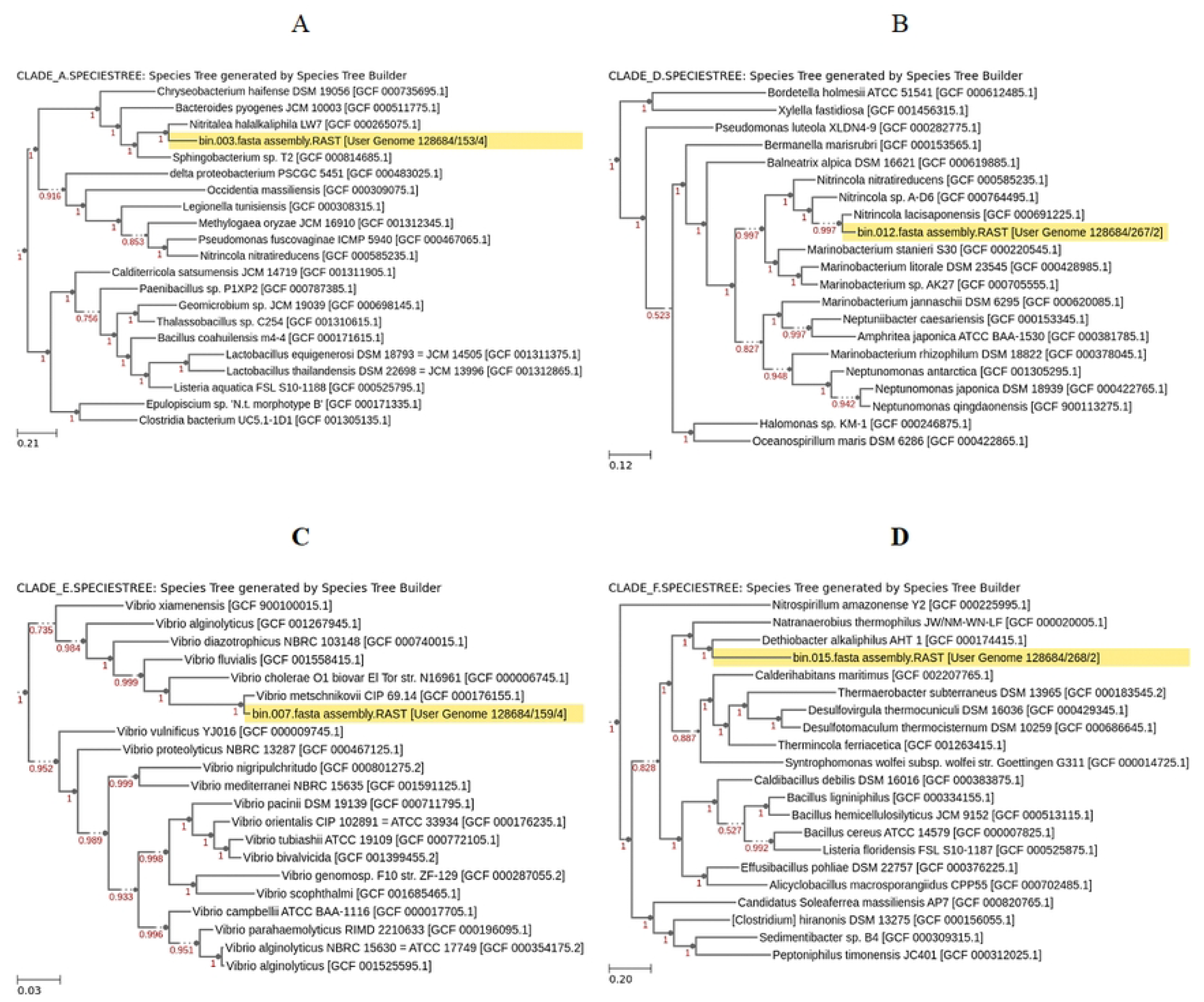
Phylogenetic placement of selected MAGs with proximal reference genomes from RefSeq. Four MAGs are shown each in its own phylogenetic tree; bin 003 (A), 012 (B), 007 (C), and 015 (D). Scale bars show branch length distance that approximates the amino acid substitution distances between the genomes in the tree, as measured from the protein multiple sequence alignments of the single-copy universal COGs found in the genomes. Red values at nodes indicate the FastTree-261 local support values for placing a split at that node, with 1 showing maximal support in the topological placement.

The 4 MAGs shown appeared to cluster closely with related organisms in the different species trees (Figure 5). MAG 003 clustered closely with *Nitritalea halalkaliphila* LW7, an alkaliphile isolated from Lonar Lake in Maharastra, India, which has an optimum pH for growth of 10.0 – 10.5 and optimum salinity of 2300 – 6500 mg/l (Jangir *et al*., 2012). Similarly, MAG 007 clusters closely with members of the *Vibrio* genus and is in all likelihood a member of this genus as well. Bacteria belonging to the *Vibrio* genus have been commonly isolated from marine as well as saline and saline-alkaline aquatic environments (Sampaio *et al*., 2022). In previous work, we discovered that the *Vibrio* is the most abundant bacterial genus in the Buhera soda pans (Mangoma *et al*., unpublished). Meanwhile, MAG 012 clustered closely to *Nitrincola lacisaponensis*, and appears to belong to the same genus (Figure 5). Genus Nitrincola is composed of different haloalkaliphilic bacterial species belonging to family *Oceanospirillaceae*, with several of its members having been isolated from soda pans and lakes (Joshi *et al*., 2020; Borsodi *et al*., 2017). Lastly, MAG 015 paired with *Dethiobacter alkaliphus* AHT 1 in the phylogenetic tree. *Dethiobacter* is an obligately haloalkaliphilic, facultatively chemolithoautotrophic, sulphidogenic, rod-shaped and anaerobic bacterium isolated from soda lake sediments and capable of growing through respiring sulphur and thiosulphate (Sorokin *et al*., 2018).

Bacterial lineages that are relatively more abundant in a microbial community are likely to be recovered as high-quality MAGs from short-read metagenomic sequences (Chivian *et al*., 2023). The greater relative abundance of these dominant lineages in the microbial community ensures sufficient read representation in the sequencing libraries, making it possible to assemble more complete genomes of these particular lineages. Due to their higher relative abundance, these leading lineages are likely to play more dominant roles in the microbiome, including contributing a larger proportion of the biomass and energy flowing through the microbiome (Chivian *et al*., 2023). Thus, the recovery of several halophilic/ haloalkaliphilic MAGs in the Buhera soda pans suggests their abundance in this environment, and may point to them playing dominant roles in this ecosystem. Thus, understanding the metabolic capacity of the microbiome is important in deciphering specific ecological roles within their natural ecosystem.

### 3.7 Functional profiling reveal possible ecological roles of MAGs

Analysing the functional profiles of the MAGs, one gets greater insight into individual organisms’ mode of life as well as major processes that are possibly active within the environment. For instance, functional analysis revealed that the MAGs recovered in this study possess a wide array of carbohydrate-metabolising pathways, with most MAGs showing versatility in the pathway used for carbohydrate metabolism (Figure 6).

**Figure 6:**
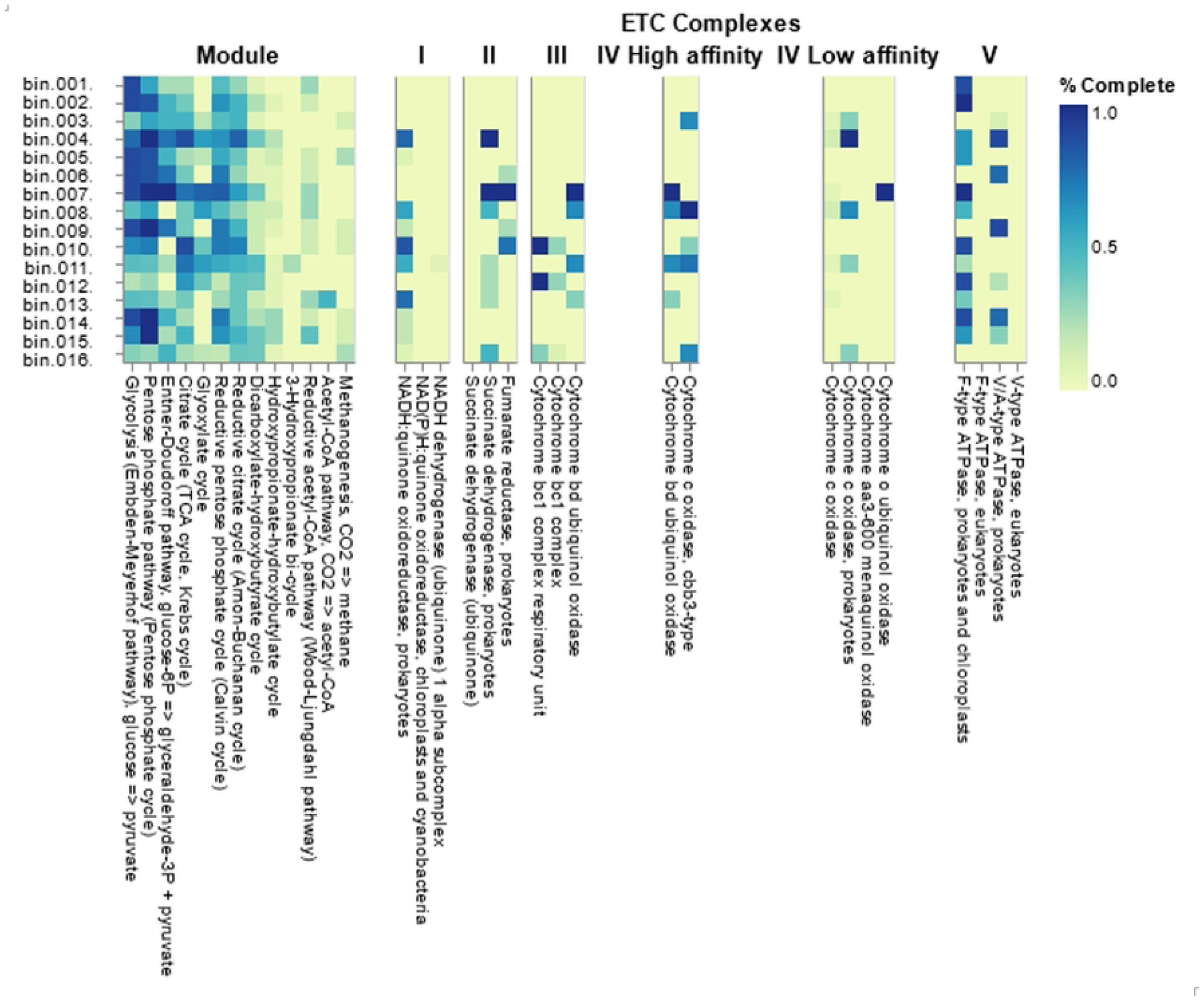
DRAM functional profiling and classification of the Buhera soda pan MAGs. The distribution of major pathways for carbohydrate metabolism and electron transport complexes within the MAG community is shown. The MAGs are shown as rows and the various metabolic processes as columns. The intensity of each cell indicates the fraction of functional marker genes (% complete) for the pathway.

Based on pathway completeness, it appeared as though a majority of the MAGs have a preference for oxidizing carbohydrates using the Embden-Meyerhof-Parnas (EMP) pathway, followed by the Pentose phosphate cycle, Entner-Doudoroff pathway and Tricarboxylic Acid (Krebs) Cycle, roughly in that order. These pathways are generally used to oxidise carbohydrates, generating electrons that are used in energy generation as well as various organic intermediates that are used in cellular biosynthesis. Most microorganisms have the ability to switch between different carbohydrate-oxidising pathways in response to changes in substrate availability or other environmental stimulus. The ability of microorganisms to switch between different energy-generating, carbohydrate-oxidising pathways ensures survival in the face of stressful conditions such as substrate depletion or the displacement of microorganisms to new environments with different carbon sources. The presence of genetic elements for the Calvin cycle in some MAGs suggest an autotrophic mode of carbon acquisition by these MAGs. Further analysis showed evidence of the use of different electron transport complexes by the Buhera soda pan microbial community. This, coupled by the presence of various oxidative processes such as the EMP pathway, Entner Doudoroff pathway and the TCA cycle, points to the presence of respiratory processes among the soda pan microorganisms. Additionally, low levels of nitrogen and sulphur metabolism were observed among the MAGs. A high level of fermentative capacity was observed, as evidenced by the many pathways for short chain fatty acid (SCFA) and alcohol conversions present among the MAGs. This suggests the use of fermentative processes as alternative energy-generating pathways by the microbial community. In fact, a lot of environmental microorganisms are facultative anaerobes and are capable of switching between respiration and fermentation as and when necessary. Further analysis of the microbial community also showed capacity to produce numerous carbohydrate-degrading or modifying enzymes.

The possible roles of selected MAGs in a number of different biogeochemical pathways was summarised and is shown in Figure 7.

**Figure 7:**
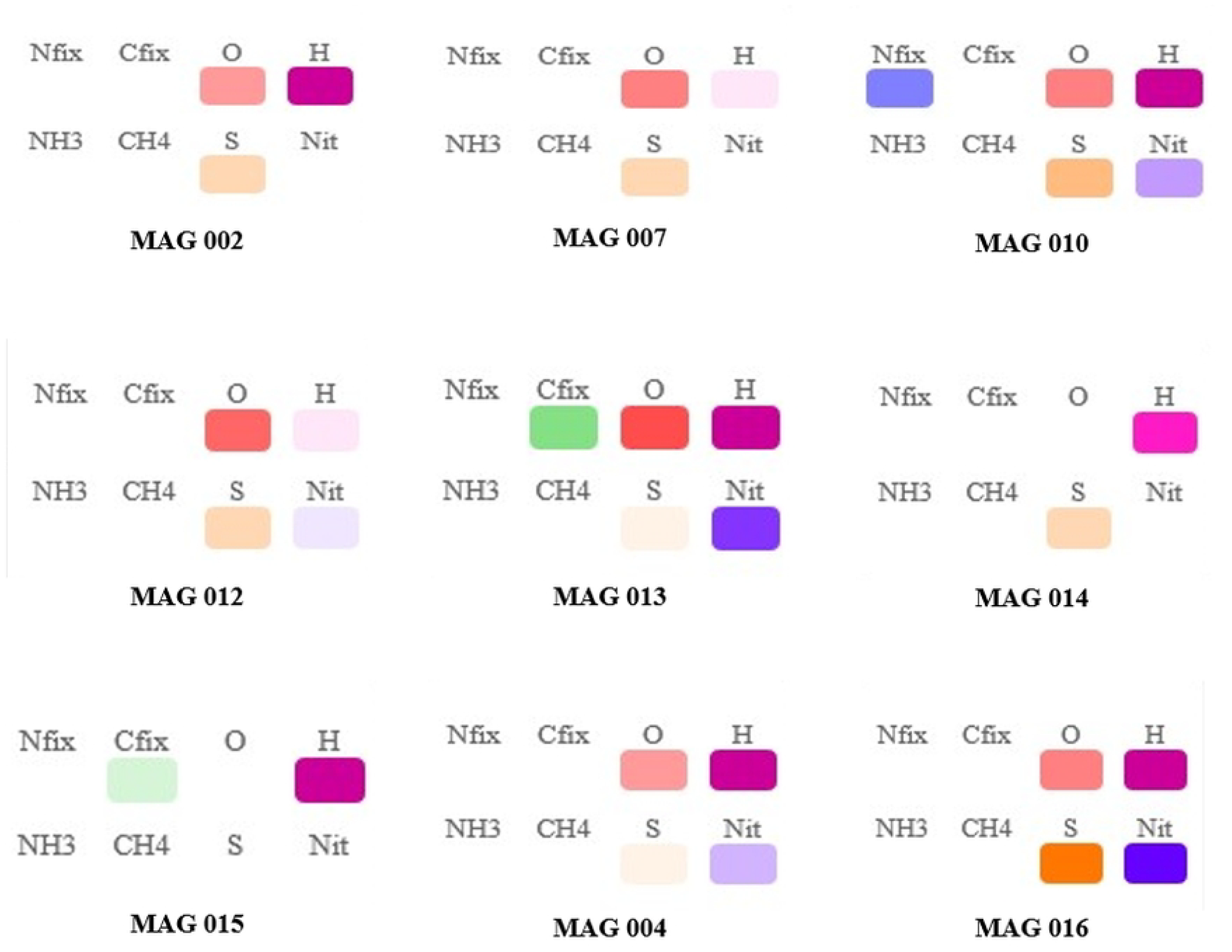
Prevalence of gene families involved in key biogeochemical transformations found using MicroTrait (https://github.com/ukaraoz/microtrait). Bioprocess categories are shown using different colours as follows: nitrogen-fixing genes (blue), carbon-fixing genes (green), oxygen-interacting genes (red), hydrogenases (magenta), ammonia-interacting genes (light blue), methane cycle genes (teal), sulfur and sulfate redox genes (orange) and nitrate redox genes (blue). A more intense box colour indicates a larger number of genes found in that category, whereas absence of a box means no genes were found for that function.

Several of the MAGs were shown to possess genes that allow for their involvement in numerous key biotransformations that include carbon and nitrogen fixation, methane cycling and sulphur/ sulphate oxidation. Only one MAG (MAG 010) was shown to be capable of nitrogen fixation, being in possession of the genes *nifD, nifK* and *nifH*. The three genes *nifD, nifK* and *nifH* encodes different components of nitrogenase, the enzyme responsible for bacterial biological nitrogen fixation. The two genes *nifD* and *nifK* encode the heterotetrameric core of bacterial nitrogenase, while *nifH* encodes the dintrogenase reductase subunit of this enzyme (Gaby and Buckley, 2014). Earlier taxonomic classification had shown MAG 010 to match closely with the uncultured bacterial genome *Campylobacteriacea* bacterium UBA1877. In addition to nitrogen fixation activity, this MAG appears to possess pathways for other crucial metabolic activities such as sulphur and sulphate reduction and nitrate reduction (Figure 7). A circular genomic map of MAG 010 confirms the presence of genes responsible both nitrogen fixation and sulphate reduction, among other key elements (Figure 8).

**Figure 8:**
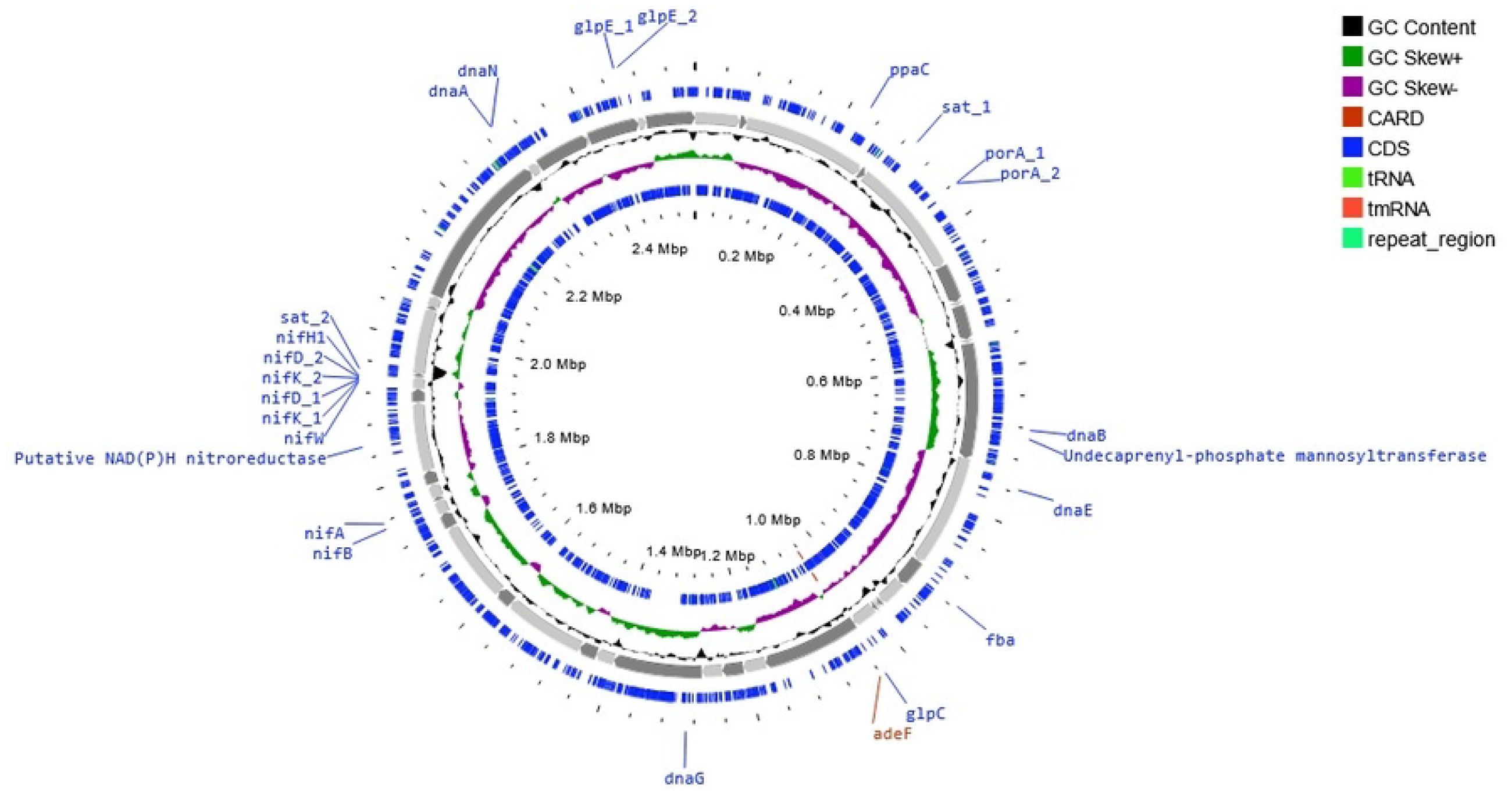
Whole genome map of MAG 010. From outside to inside: coding sequences (CDS) located on the forward strand (blue), sequence backbone made up of 46 contigs joined end-to-end (grey), GC content (black), GC skew (purple when GC skew < 0, or olive when GC skew > 0), coding sequences (cds) located on the reverse strand (blue).

As shown, MAG 010 possesses two clusters of genes (possibly operons) supporting nitrogen fixation. The first cluster contains the the *nifA* and *nifB* genes and is located on the reverse strand of contig 29 (NODE_476_length_21671_cov_6.024796) in the MAG map, and the second cluster, located on the reverse strand of contig 35 (NODE_549_length_18964_cov_5.559099), contains the *nifD, nifH, nifK* and *nifW* genes. The *nifA* gene encodes a regulator protein, which when expressed (under conditions of low fixed nitrogen availability) activates the transcription of the rest of the *nif* gene complex, leading to their expression and eventual nitrogen fixation. In addition to the nitrogen fixation complex, genes involved in sulphur reduction can also be located on the map, for example the *sat1* and *sat2* genes which encode the enzyme sulphate adenyltransferase (EC 2.7.7.4) which catalyses the adenylation of sulphate using ATP to synthesise adenylyl sulphate and pyrophosphate, the latter which can be hydrolysed by the *ppaC* (pyrophosphatase) gene which was also located on the forward strand of contig 11 (NODE_11_length_182382_cov_6.080246). These enzymes play a key function in both assimilatory sulphur reduction and dissimilatory sulphur oxidation, and are hence key to the sulphur cycle. The presence of the *fba* gene which encodes the enzyme Fructose 1,6 bisphosphate aldolase suggests a possible mechanism of carbohydrate metabolism through the glycolytic pathway by this MAG. Other genes shown on the map include an antibiotic resistance gene *adeF* which encodes a drug efflux pump, and genes encoding some subunits of the DNA polymerase enzyme (*dnaA, dnaB, dnaE, dnaG* and *dnaN*) involved in DNA replication.

On the other hand, only 2 of the 16 MAGs were shown to be autotrophic, with one being a photoautotroph (MAG 13) and the other one a chemolithoautotroph (MAG 15). MAG 13 was classified under unclassified bacterial class *Anaerolineae* under the photoautotrophic bacterial phylum *Chloroflexi*, while MAG 15 was shown to be an unclassified *Dethiobacteria* under phylum *Firmicutes*. The presence of these primary producers in the ecosystem is important as they contribute to the production of organic matter required by the heterotrophic fraction of the soda pan microbiome. In addition to these annotations, the metabolic capacity of the different MAGs and their potential contribution to important biogeochemical transformations was summarized and presented in Figure 9.

**Figure 9:**
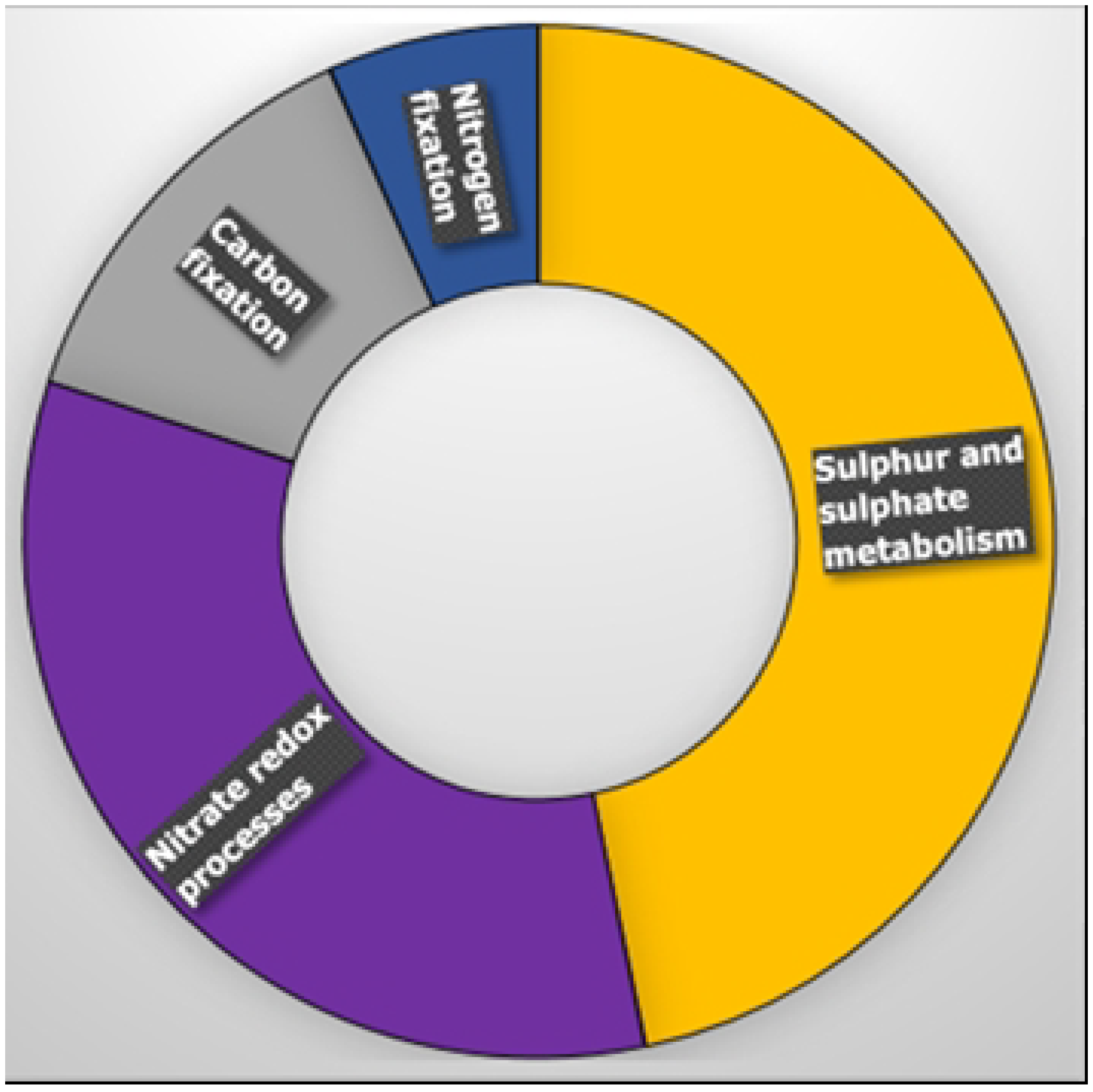
Sunburst chart showing distribution of genes supporting various biogeochemical processes among the MAGs.

The majority of MAGs were shown to possess pathways for participation in two major processes, sulphur and sulphate metabolism and nitrate oxidation and reduction processes. The greater prevalence of sulphur and sulphate metabolism supports the hypothesis that a significant proportion of organisms in this ecosystem respire anaerobically using sulphur and/ or sulphate as terminal electron acceptors. These findings are supported by our earlier observations showing the dominance of sulphur compounds in the Buhera soda pans. Overall, the results of the functional analysis show that different biogeochemical processes are active within the Buhera soda pans.

## Conclusion

In this study, we report on the recovery of and annotation of 16 microbial genomes, all belonging to five phyla within the bacteria domain, from a previously unexplored and uncharacterised extreme natural environment. Six of the reconstructed genomes could not match any cultured reference strains, suggesting that the Buhera soda pans may harbour a large number of previously uncultured bacteria. Most of the MAGs were of medium-to-high-quality, harbouring sufficient genetic information to enable their detailed annotation. Five of the sixteen MAGs belonged to halophilic/ haloalkaliphilic genera, suggesting a possible dominance of these salt- and alkali-loving extremophiles in the Buhera soda pans. Functional annotation of the MAGs revealed a wide array of metabolic pathways among the MAGs, with key processes being sulphur and sulphate metabolism, nitrate reduction and carbon and nitrogen fixation. More work is needed in order to allow us to learn more about the composition and biotechnological potential of this complex microbial habitat.

## Conflict of interest

The authors have no conflict of interest to declare.

## Funding

This work was partly funded by the Research and Development Board of the National University of Science and Technology, Bulawayo, Zimbabwe. Funds for metagenomic sequencing was provided by the Botswana International University of Science and Technology Research Office.

## Acknowledgements

This work was carried out in the Biochemistry, Microbiology and Biotechnology laboratories at the National University of Science and Technology, Bulawayo, Zimbabwe. The authors would like the acknowledge the Gombahari village leadership and local community for granting them access to the soda pans.

## Notes

### Competing Interest Statement

The authors have declared no competing interest.

## References

Alcock, B.P., Huynh, W., Chalil, R., Smith, K.W., Raphenya, A.R., Wlodarski, M.A., Edalatmand, A., Petkau, A., Syed, S.A., Tsang, K.K., Baker, S.J.C., Dave, M., McCarthy, M.C., Mukiri, K.M., Nasir, J.A., Golbon, B., Imtiaz, H., Jiang, X., Kaur, K., Kwong, M., Liang, Z.C., Niu, K.C., Shan, P., Yang, J.Y.J., Gray, K.L., Hoad, G.R., Jia, B., Bhando, T., Carfrae, L.A., Farha, M.A., French, S., Gordzevich, R., Rachwalski, K., Tu, M.M., Bordeleau, E., Dooley, D., Griffiths, E., Zubyk, H.L., Brown, E.D., Maguire, F., Beiko, R.G., Hsiao, W.W.L., Brinkman, F.S.L., Van Domselaar, G. and McArthur, A.G. 2023. CARD 2023: expanded curation, support for machine learning, and resistome prediction at the Comprehensive Antibiotic Resistance Database. Nucleic Acids Research. 10.1093/nar/gkac920.

Alneberg, J. 2018. Bioinformatic Methods in Metagenomics. Doctoral Thesis, KTH Royal Institute of Technology Engineering Sciences in Chemistry, Biotechnology and Health Department of Gene Technology Science for Life Laboratory SE-171 65 Solna, Sweden.

Boros, E. and Kolpakova, M. 2018. A review of the defining chemical properties of soda lakes and pans: An assessment on a large geographic scale of Eurasian inland saline surface waters. PLoS ONE 13(8): e0202205. 10.1371/journal.pone.0202205.

Borsodi, A.K., Korponai, K., Schumann, P., Sproer, C., Felfordi, T., Marialigeti, K., Szilikovacs, T. and Toth, E. 2017. Nitrincola alkalilacustris sp. nov. and Nitrincola schmidtii sp. nov., alkaliphilic bacteria isolated from soda pans, and emended description of the genus Nitrincola. International Journal of Systematic and Evolutionary Microbiology, 67(12), pp. 5159–5164. 10.1099/ijsem.0.002437.

Bowers, R.M., Kyrpides, N.C., Stepanauskas, R., Harmon-Smith, M., Doud, D., Reddy, T.B.K., Schulz F., Jarett, J., Rivers, A.R., Eloe-Fadrosh, E.A., Tringe, S.G., Ivanova, N.N., Copeland, A., Clum, A., Becraft, E.D., Malmstrom, R.R., Birren, B., Podar, M., Bork, P., Weinstock, G.M., Garrity, G.M., Dodsworth, J.A., Yooseph, S., Sutton, S., Glöckner, F.O., Gilbert, G.A., Nelson, W.C., Hallam, S.J., Jungbluth, S.P., Ettema, T.J.G., Tighe, S., Konstantinidis, K.T., Liu, W-T., Baker, B.J., Rattei, T., Eisen, J.A., Hedlund, B., McMahon, K.D., Fierer, N., Knight, R., Finn, R., Cochrane, G., Karsch-Mizrachi, I., Tyson, G.W., Rinke, C. The Genome Standards Consortium, Lapidus, A., Meyer, F., Yilmaz, P., Parks, D.H., Eren, A.M., Schrim, L., Banfield, J.F., Hugenholtz, P. and Woyke. T. 2017. Minimum information about a single amplified genome (MISAG) and a metagenome-assembled genome (MIMAG) of bacteria and Archaea. Nature Biotechnology 35(8), pp. 725–731. 10.1038/nbt.3893.

Bryanskaya, A.V., Shipova, A.A., Rozanov, A.S., Kolpakova, O.A., Lazareva, E.V., Uvarova, Y.E., Efimov, V.M., Zhmodik, S.M., Taran, O.P., Goryachkovskaya, T.N., Peltek, S.E. 2022. Diversity and Metabolism of Microbial Communities in a Hypersaline Lake along a Geochemical Gradient. Biology 11(605). 10.3390/biology11040605.

Chivian, D., Jungbluth, S.P., Dehal, P.S., Wood-Charlson, E.M., Canon, R.S., Allen, B.H., Clark, M.M., Gu, T., Land, M.L., Price, G.A., Riehl, W.J., Sneddon, M.W., Sutormin, R., Zhang, Q., Cottingham, R.W., Henry, C.S. and Arkin, A.P. 2022. Metagenome-assembled genome extraction and analysis from microbiomes using KBase. Nature Protocols, 18(1), pp.208–238. 10.1038/s41596-022-00747-x.

Choure, K., Parsai, S., Kotoky, R., Srivastava, A., Tilwari, A., Rai, P.K., Sharma, A. and Pandey, P. 2021. Comparative Metagenomic Analysis of Two Alkaline Hot Springs of Madhya Pradesh, India and Deciphering the Extremophiles for Industrial Enzymes. Frontiers in Genetics, 12. 10.3389/fgene.2021.643423.

Couvin, D., Bernheim, A., Toffano-Nioche, C., Touchon, M., Michalik, J., Néron, B., Rocha, E.P.C., Vergnaud, G., Gautheret, D. and Pourcel, C. 2018. CRISPRCasFinder, an update of CRISRFinder, includes a portable version, enhanced performance and integrates search for Cas proteins. Nucleic Acids Research. 10.1093/nar/gky425.

Delcher, A.L., Bratke, K.A., Powers, E.C. and Salzberg, S.L. 2007. Identifying bacterial genes and endosymbiont DNA with Glimmer. Bioinformatics, 23(6), pp.673–679. 10.1093/bioinformatics/btm009.

Felföldi, T. 2020. Microbial communities of soda lakes and pans in the Carpathian Basin: a review. Biologia Futura 71:393–404. 10.1007/s42977-020-00034-4.

Gaby, J.C. and Buckley, D.H. 2014. A comprehensive aligned nifh gene database: A multipurpose tool for studies of nitrogen-fixing bacteria’, Database, 2014. 10.1093/database/bau001.

Govil, T., Sharma, W., Salem, D.R. and Sani, R.K. 2021. Multi-Omics Approaches for Extremophilic Microbial, Genetic, and Metabolic Diversity, In: “Extreme Environments”. 1st Edition. Imprint CRC Press. eBook ISBN 9780429343452.

Grötzinger, S.W., Karan, R., Strillinger, E., Bader, S., Frank, A., Rowaihi, A., Anastassja Akal, Wiebke Wackerow, Archer, J., Magnus Rueping, Weuster-Botz, D., Groll, M., Eppinger, J. and Arold, S.T. 2018. Identification and Experimental Characterization of an Extremophilic Brine Pool Alcohol Dehydrogenase from Single Amplified Genomes. ACS Chemical Biology, 13(1), pp.161–170. 10.1021/acschembio.7b00792.

Guan, T.W., Lin, Y.J., Ou, M.Y. and Chen, K B. 2020. Isolation and diversity of sediment bacteria in the hypersaline aiding lake, China. PLoS ONE 15(7): e0236006. 10.1371/journal.pone.0236006.

He, Y., He, L., Wang, Z., Liang, T., Sun, S. and Liu, X. 2022. Salinity Shapes the Microbial Communities in Surface Sediments of Salt Lakes on the Tibetan Plateau, China. Water, 14, 4043. 10.3390/w14244043.

Hyatt, D., Chen, G.-L., LoCascio, P.F., Land, M.L., Larimer, F.W. and Hauser, L.J. (2010). Prodigal: prokaryotic gene recognition and translation initiation site identification. BMC Bioinformatics, 11(1). 10.1186/1471-2105-11-119.

Jangir, P.K., Singh, A., Shivaji, S. and Sharma, R. 2012. Genome sequence of the alkaliphilic bacterium Nitritalea halalkaliphila type strain LW7, isolated from Lonar Lake, India. Journal of Bacteriology, 194(20), pp. 5688–5689. 10.1128/jb.01302-12.

Joshi, A., Thite, S., Dhotre, D., Moorthy, M., Joseph, N., Ramana, V.V. and Shouche, Y. 2020. Nitrincola tapanii sp. nov., a novel alkaliphilic bacterium from an Indian Soda Lake. International Journal of Systematic and Evolutionary Microbiology, 70(2), pp. 1106–1111. 10.1099/ijsem.0.003883.

Land, M., Hauser, L., Jun, S-R., Nookaew, I., Leuze, M.R., Ahn, T-H., Karpinets, T., Lund, O., Kora, G., Wassenaar, T., Proudel, S. and Ussery, D.W. 2015. Insights from 20 years of bacterial genome sequencing. Functional & Integrative Genomics 15(2), pp. 141–161. 10.1007/s10142-015-0433-4.

Mangoma, N., Siwela, A.H., Zhou, N. and Ncube, T. Unpublished. Metagenomic analysis of the microbial and functional diversity of the Manicaland Soda pans in Zimbabwe.

Nyakeri, E.M., Mwirichia, R. and Boga, H. 2018. Isolation and characterization of enzyme-producing bacteria from Lake Magadi, an extreme soda lake in Kenya. Journal of Microbiology and Experimentation, 6(2). 10.15406/jmen.2018.06.00189.

Omeroglu, E.E., Sudagidan, M., Yurt, M.N.Z., Tasbasi, B.B., Acar, E.E. and Ozalp, V.C. 2021. Microbial community of soda Lake Van as obtained from direct and enriched water, sediment and fish samples. Nature Scientific Reports. 10.1038/s41598-021-97980-3

Parks, D.H., Imelfort, M., Skennerton, C.T., Hugenholtz, P. and Tyson, G.W. 2014. Checkm: Assessing the quality of microbial genomes recovered from isolates, single cells, and Metagenomes. Genome Research, 25: 1043–1055.

Sampaio, A., Silva, V., Poeta, P. and Aonofriesei, F. 2022. Vibrio spp.: Life Strategies, ERcology, and Risks in a Changing Enviornment. Diversity 14(2). 10.3390/d14020097.

Seemann, T. 2014. Prokka: rapid prokaryotic genome annotation. Bioinformatics 10.1093/bioinformatics/btu153.

Singh, N.K., Wood, J.M., Patane, J., Moura, L.M.S., Lombardino, J., Setubal, J.C. and Venkateswaran, K. 2023. Characterization of metagenome-assembled genomes from the International Space Station. Microbiome 11(1). 10.1186/s40168-023-01545-7.

Sorokin, D.Y., Muntyan, M.S., Toshchakov, S.V., Korzhenkov, A. and Kublanov, I.V. 2018. Phenotypic and Genomic Properties of a Novel Deep-Lineage Haloalkaliphilic Member of the Phylum Balneolaeota From Soda Lakes Possessing NaC-Translocating Proteorhodopsin. Frontiers in Microbiology. 9:2672. 10.3389/fmicb.2018.02672.

Sysoev, M., Grötzinger, S.W., Renn, D., Eppinger, J., Rueping, M. and Karan, R. 2021. Bioprospecting of Novel Extremozymes From Prokaryotes—The Advent of Culture-Independent Methods. Frontiers in Microbiology 12:630013. 10.3389/fmicb.2021.630013.

https://narrative.kbase.us/#catalog/apps/kb_gtdbtk/run_kb_gtdbtk

https://github.com/ukaraoz/microtrait

